# Regulation of cell cycle by the novel GATA1/TAL1/Sphingomyelin Synthase 1 (*SGMS1*) transcriptional axis. Implications for anti-leukemic strategies

**DOI:** 10.1101/2025.01.19.633812

**Authors:** Yasharah Raza, Sitapriya Moorthi, Gui-Qin Yu, Sam Chiappone, Caterina Vacchi-Suzzi, Chiara Luberto

## Abstract

Sphingomyelin Synthase 1 (gene name: *SGMS1*) participates in regulation of sphingolipid levels by synthesizing sphingomyelin from ceramide and phosphatidylcholine. Evidence have supported SGMS1’s functions in regulating proliferation, cell cycle, cell death and migration. While its functions have begun to be explored, very little is known about upstream regulators. Here, we demonstrate that *SGMS1* is a direct gene target of the GATA1-TAL1 transcriptional complex in K562 erythroleukemic cells. A predicted GATA1 consensus DNA binding sequence was identified with in a newly characterized alternative *SGMS1* promoter (TSS-7 promoter) and ChIP analysis confirmed GATA1 occupancy on the promoter. Down-regulation of GATA1 in K562 cells significantly decreased SGMS1 expression and enzymatic activity, and ChIP-Seq analysis from ENCODE showed colocalized peaks for GATA1 and TAL1 (a well-established GATA1 DNA binding partner) on the *SGMS1* gene. Analysis of publicly available datasets shows that elevated *GATA1, TAL1* and *SGMS1* expression not only clusters GATA1 positive chronic myelogenous leukemia cells (like K562), but also selectively identifies acute erythrocytic and megakaryocytic leukemias (M6 and M7 AML, respectively). Microarray gene expression analysis after down-regulation of *SGMS1* in M6 AML Hel cells revealed alteration of genes regulating G2/M check point and mitotic spindle formation. This phenotype was functionally confirmed by the significant delay in G2/M cell cycle progression of cells with *SGMS1* downregulation and sensitization to the clinically relevant anti-mitotic agent, Taxol. Altogether, these results identify *SGMS1* as a novel target of GATA1/TAL1 transcriptional complex and they support a role for the GATA1/TAL1/SGMS1 axis in regulating transit through G2/M. Importantly, results also point to combination of anti-mitotic agents and inhibition of SGMS1 as a potential novel therapeutic approach against the aggressive and resilient M6 AMLs.

**KEY POINTS:**

- The Sphingomyelin Synthase 1 gene (*SGMS1*) is a novel direct target of GATA1 and TAL1.
- High *SGMS1* levels are associated with high *GATA1/TAL1* expression and regulate cell cycle progression through the G2/M checkpoint in GATA1^+^ erythroleukemic Acute Myeloid Leukemia Hel cells.
- High *SGMS1* is associated with lower probability of survival of patients with Acute Myeloid Leukemia and down-regulation of SGMS1 co-operates with microtubule targeting agents to induce cytotoxicity in GATA1 positive AML Hel cells.

## INTRODUCTION

Sphingomyelin synthase (SMS) represents a class of enzymes that synthesizes sphingomyelin (SM), a major component of the outer leaflet of the plasma membrane and homeostatic modulator of lipid microdomains(1, 2). SMS catalyzes this reaction using ceramide (Cer) and phosphatidylcholine to generate SM and diacylglycerol (DAG)(2–4). Ceramide, the central bioactive sphingolipid, exerts mostly anti-proliferative and pro-inflammatory effect on cells(5–7), while DAG has mitogenic effects and pro-secretory functions *via* recruitment to the membrane and/or activation of various protein kinases (such as PKCs, PKDs) and Ras (indirectly)(3, 4, 8–11). Thus, SMS, by regulating SM, DAG and Cer, is uniquely positioned to modulate important cellular functions through SM-dependent and SM-independent mechanisms(12–20). In mammalian cells, SMS activity is carried out by two iso-proteins, SMS1 and SMS2, encoded by *SGMS1* and *SGMS2*, respectively(21, 22).

At the cellular level, SMS1 and SMS2 impact proliferation, apoptosis, inflammation and secretion in ways that depend on the cell context (23–29). For example, in the murine neuroblast cell line Neuro-2a, inhibition of SMS1 activity is sufficient to reduce proliferation and causes cell cycle arrest(30) while in breast cancer MDA-MB-231 cells, its overexpression inhibits proliferation and migration(31). On the other hand, in breast cancer cells MCF-7 and MDA-MB-231, SMS2 supports proliferation and migration through inhibition of apoptosis(32) and by promoting epithelial-to-mesenchymal transition (EMT), respectively (32). While existing links between SMSs and oncogenic processes suggest their potential usefulness as therapeutic targets depending on the malignancy, mechanisms governing *SMSs* gene expression are largely unknown.

Our previous work demonstrated upregulation of *SGMS1*/SMS1 expression and activity in the Chronic Myelogenous Leukemia (CML) K562 cells, where we linked enhanced SMS1 activity to the cells’ proliferative capacity (33). Furthermore, we showed that up-regulation of *SGMS1* occurs *via* increased transcription from a newly identified alternative transcription start site (TSS-7) regulated by a previously unknown promoter juxtaposed to and spanning the 5’ end of *SGMS1’s* Exon 7 (TSS-7 promoter)(34). While the only known mechanism of transcriptional regulation of *SGMS1* is *via* its repression by FOXO1 in dental pulp stem cells (35), understanding the molecular mechanisms leading to *SGMS1* transcription (from canonical or alternative TSSs) could provide critical insight into other biological processes where *SGMS1* plays a role.

Herein, starting from the analysis of the TSS-7 promoter and chromatin immunoprecipitation, we uncover positive regulation of *SGMS1* by the transcription factor GATA1 in complex with TAL1 in K562 cells. Investigating this novel regulatory link, we identified acute erythroleukemia and acute megakaryocytic leukemia (M6 and M7 AML) as types of leukemias that exhibit a strong positive *GATA1, TAL1, SGMS1* correlation. Functionally, in erythroleukemic M6 AML cells, we linked the high *SGMS1* expression with regulation of cell cycle progression through the G2/M checkpoint, we leveraged this phenotype to promote ER stress and, by targeting SM synthesis, we sensitized these aggressive leukemic cells to the anti-mitotic agent, Taxol.

## MATERIALS AND METHODS

### Cell lines

K562 and HL-60 cell lines were from ATCC (Rockville, MD). Hel, LAMA-84 and JURL-MK-1 were from DSMZ (Brunswick, Germany). Molm-16, K562, LAMA-84 and JURL-MK-1 were grown in RPMI-1640 with 10% Heat Inactivated-Fetal Bovine Serum (HI-FBS). HL-60 were grown in RPMI with 20% HI-FBS. Growth medium was supplemented with 100 U/mL Penicillin and 100 μg/mL Streptomycin. RPMI, HI-FBS, Penicillin and Streptomycin were from Life Technologies Corporation (Carlsbad, CA). Cells were maintained at 37°C and 5% CO_2_ in a humidified incubator and routinely checked for mycoplasma contamination with the Lonza mycoAlert detection kit (Morristown, NJ, USA). Hel cells were treated with Taxol (Sigma Aldrich; St. Louis, MO, USA) or HPA-12 (Cayman Chemical Company Incorporated; Ann Arbor, MI; 28350) as indicated in the respective figure legend.

### Plasmid constructs

#### *SGMS1* TSS-7-5’ progressive promoter deletion constructs

Promoter deletion constructs were generated from the full-length TSS-7 promoter luciferase construct (**Supplementary Table 1**). PCR was performed using forward primers designed to progressively exclude small stretches of the TSS-7 promoter, starting at the 5’ end **(Supplementary Table 2)**. PCR amplicons were cloned into the TOPO-TA vector system (Thermo Fisher Scientific, Waltham, MA) followed by sub-cloning into pGL3-basic vector (Promega, Madison, WI, USA) as previously described (34). Construct integrity was validated by sequencing.

#### *SGMS1* TSS-7 Region I, Region II and Region I +II promoter deletion constructs

Promoter constructs lacking Region I, Region II or both Region I and II were generated by Life Technologies Corporation (Carlsbad, CA) (sequences in **Supplementary Table 3)**. Constructs were sub-cloned into pGL3-basic vector (Promega, Madison, WI, USA) as previously described(34).

### Transient transfections

Transient transfections were performed using the Neon Electroporation system (Life Technologies Corporation, Carlsbad, CA). Unless otherwise noted, each transfection mix contained: 2×10^6^ cells (exponentially growing) in 100 μL Buffer-R and 20 μL of endotoxin free plasmid or siRNA diluted in buffer R. After electroporation, cells were transferred to a 6-wells plate with 2 mL complete pre-warmed media. After 1 hour recovery, cells were diluted to a concentration of 0.2×10^6^ cells/mL (unless otherwise stated). Electroporation parameters are: 1000V, 50 ms, 1 pulse.

#### Transient co-transfection of TSS-7 promoter luciferase deletion constructs and β-galactosidase reporter

Five μg of pGL3 basic vector or pGL3 basic-TSS-7 promoter constructs and 5μg of CMV-β-galactosidase reporter construct (Promega, Madison, WI) were co-transfected into cells. Cells were incubated in recovery conditions (1×10^6^ cells/mL) and collected after 16 hours.

#### Transient co-transfection of TSS-7 promoter luciferase construct and β-galactosidase after *GATA1* siRNA mediated knock-down

Cells were transfected with *GATA1* siRNA (as indicated below) and diluted to 0.2×10^6^ cells/mL. After 32 hours, cells were transfected with luciferase and β-galactosidase constructs (as above) and collected for luciferase and β-galactosidase activity 16 hours after plasmid transfection.

#### Transient siRNA transfection of K562 and Hel cells

For human *GATA1* knockdown, GATA1 siRNA was sc-29330 from Santa Cruz Biotechnologies (Santa Cruz, CA) and control siRNA was Allstar negative control from Qiagen (Germantown, MD) (20nM of siRNA used). For human *GATA2* knockdown, siRNA was GATA2 Silencer Select S5598 from Ambion (Austin, TX) and Ambion Silencer Select Negative Control #1 (12nM of siRNA used). The protocol was optimized for siRNA concentration, transfection and post-transfection conditions in preliminary experiments (data not shown). SiRNA concentrations are calculated considering 10 ml of final volume post-electroporation.

For microarray analysis after *SGMS1* knockdown in Hel cells, 2×10^6^ cells were transfected using 30nM siRNA as previously described (33); immediately after transfection, cells were diluted in 4ml of growth medium (siRNA concentrations are calculated considering 4 ml of final volume post-electroporation). To achieve lasting knockdown, three siRNA transfections were repeated every other day and, after 48 hours from the third transfection, cells were collected for RNA extraction.

#### Lentiviral knockdown of *SGMS1*

For lentiviral experiments, cells were transduced with either non-targeting control particles (shRNA-NTCP; SHC002V) or Mission lentiviral transduction particles (both from Sigma-Aldrich) that contained the pLKO.1-puro vector including the *SGMS1* targeting sequence, 5’-CCG GCC AAC TGC GAA GAA TAA TGA ACT CGA GTT CAT TAT TCT TCG CAG TTG GTT TTT TG-3’ (sh-RNA-SMS1; SHCLNV, TRCN0000133921). During transduction, 3×10^4^ cells/ml re-suspended in RPMI-1640 containing 10% FBS and 10 μg/ml of Hexabromide (Sigma) were infected at a Multiplicity of Infection (MOI) of 10 with the appropriate lenti-viral particles. Cells were incubated at 37°C, 5% CO_2_ and 3 days post-transduction, 10 μg/ml of puromycin (Sigma) was added to the media to select for pLK0.1-puro positive clones and re-added every 3-4 days to maintain selection. The concentration of puromycin needed to select for vector expressing clones was based on a kill-curve analysis showing 70-80% cell death after 3 days in non-transduced cells (data not shown). SMS1 down-regulation was confirmed by quantitative real time PCR. Mono-clones were isolated by diluting culture and seeding single cells into a 96-well plate.

### Western Blotting

Twenty μg of lysates prepared in 0.5% SDS (Sodium Dodecyl Sulphate) (on ice) and sonicated or 60μg of total membranes (for SMS1) (33, 34, 36) were run in SDS-PAGE and blotted with primary and secondary antibodies (**Supplementary Table 4).** Quantification was performed using Image J (National Institute of Health, Bethesda, MD) and Adobe Photoshop. Statistical significance was determined by paired t-test.

### Chromatin Immunoprecipitation (ChIP)

ChIP assay was performed from 20-40×10^6^ cells using the Magna ChIP–G-immunoprecipitation kit according to the protocol (EMD Millipore, MA). Briefly, after cross-linking cells (final formaldehyde concentration 1%) at room temperature and two washes with ice cold 1x PBS, cells were lysed on ice, nuclei isolated by centrifugation and lysed with appropriate buffer. Nuclear lysate was processed by sonication (10% output, 20s x 6 cycles with 50s rest), after standardization for sonication and shearing of cross-linked chromatin. Sonicated chromatin was centrifugated and diluted in appropriate buffer. Five μL input chromatin (corresponding to 1%) was aliquoted and stored at 4°C while the rest was pre-cleared with 20μL protein G beads for 1 hour at 4°C. After centrifugation, supernatant was incubated with 6μg of IgG (purified rabbit monoclonal, ab172730, Abcam, Cambridge, MA) or GATA1 antibody (purified rabbit monoclonal, ab181544, Abcam, Cambridge, MA). Immunoprecipitation was standardized to allow for maximum pull-down and enrichment of DNA bound-GATA1. After following the manufacturer’s protocol, eluted DNA was probed by qRT-PCR using ChIP primers **(Supplementary Table 5).**

### RNA extraction and cDNA synthesis

Total RNA was isolated and cDNA was synthesized as previously described(34).

### Quantitative Real-Time PCR (qRT-PCR)

qRT-PCR was performed using SYBR green mixture (Bio-Rad, Hercules, CA) as previously indicated (34). Results were normalized to internal control gene, β-actin or 18SRNA, and amplification efficiency. Mean of normalized expression (MNE)(37) was used for analysis. For some genes, Taqman probes (from Thermo Fisher Scientific, Waltham, MA) were utilized with iTaq Universal Supermix (Bio-Rad, Hercules, CA, USA) according to manufacturer’s instructions. Primers’ sequences and Taqman probes’ catalogue numbers are provided in **Supplementary Table 5**.

### SMS Activity Assay

SMS activity was performed as previously described with modifications (3). Three to four million cells were lysed by tip sonication (11% power for 20 seconds) in 150 μl of homogenization buffer (250 mM Tris, 50 mM EDTA pH 7.4, Pierce Halt phosphatase inhibitor, Pierce Halt protease inhibitor, 5 mM PMSF). Unbroken cells were pelleted by centrifugation. Supernatant was collected and assayed for protein concentration using the Bio-Rad protein determination assay (Bio-Rad, Hercules, CA). SMS activity was assayed on 50μg proteins as previously described (3).

### GATA1 and TAL1 ChipSeq data analysis and RNASeq for TAL1 siRNA

ChIP-Seq and RNA-Seq data for *SGMS1* in K562 cells were mined from public datasets. ChIP Seq data were from the Michael Snyder Lab (Stanford) and the accession and experiment codes are as follows: GATA (ENCFF000YNI; ENCSR000EFT); TAL1 (ENCFF509LKA; ENCSR000EHB). Data were visualized using Integrative Genomics Viewer (38, 39). RNAseq data was obtained from the Encode Project. Datasets used include ENCSR336ZWX (*TAL1* siRNA knockdown followed by RNA-seq) and ENCSR641CMW (control siRNA knockdown followed by RNA-seq).

### Correlation Analysis

*Pearson Correlation Analysis of SGMS1 in CML and AML cell lines.* DepMap portal was utilized to derive gene correlations and corresponding q values between *SGMS1* and various genes (as indicated in the Figures) in CML and AML cell lines. Expression Public 24Q2 from the Cancer Cell Line Encyclopedia (CCLE) was the reference database for the analysis. *Pearson Correlation Analysis of SGMS1 in AML patient samples.* Using the TCGA database (LAML), AML patients were stratified into GATA1+ and GATA1-AML cohorts based on the expression of GATA1 in erythroleukemia/megakaryoblastic leukemia (M6/M7 AML in the French-American-British classification), which are widely considered to be GATA1 expressing leukemias. Those patients that had equal or higher *GATA1* expression to that of the lowest M6/M7 patient sample were considered GATA1+. In these RNA-Seq samples, the Pearson correlation was determined between the expression of every gene and the *SGMS1* gene, to find *SGMS1* co-expressed genes. This Pearson correlation was interrogated for known GATA1 binding partners. Significance was calculated based on the T-statistic corresponding to each Pearson correlation.

### Microarray analysis

mRNA from 4 independent experiments of *SGMS1* siRNA-mediated knock down in Hel cells was utilized. Samples had a RIN number ≥9.2. One hundred fifty ng of total RNA were processed for whole transcriptome expression analysis, according to the manufacturer’s protocol (GeneChip™ WT PLUS Reagent Kit from Applied Biosystems). A cocktail containing 1.6 µg of fragmented and labeled single stranded DNA was hybridized to Clariom S Human Arrays (Applied Biosystems) for 16hrs at 45°C at 60 RPMs in a Genechip Hybridization Oven 640. The arrays were washed and stained in a GeneChip Fluidics 450 station and scanned in a GeneChip Scanner 3000 G7 controlled by GeneChip Command Console v 4.3.3.1616 software. Data analysis was completed using Applied BioSystems Transcriptome Analysis Console software v4.0.2.15. For Gene Set Enrichment Analysis (GSEA) raw data from .CEL files were extracted and RMA normalized using the oligo package in R. Normalized data were analyzed by using GSEA java-based executable, available from Broad Institute (http://www.gsea-msigdb.org/gsea/index.jsp). For GSEA analysis, 10,000 permutations and a random seed were used; permutations were of the genes in the gene sets. The HALLMARK database for the gene sets was used.

### Cell Cycle Arrest and Analysis

Hel cells that had stable *SGMS1* downregulation via shRNA (described above) were first treated with 100 ng/mL Nocodazole (Sigma, MO) for 12 hours at a cell concentration of 0.2×10^6^ cells/mL to induce G2/M arrest; they were then released from the arrest after a wash with 1x PBS and re-suspended in the same original volume of media containing 10% FBS. Cells were collected immediately after re-suspension for the zero hour (0h) time-point and after two hours (2h) for cell cycle analysis (the 2h time point was selected as it offered the highest percentage of cells transitioning in G1 with the least amount of cells advancing further). Cells were collected by centrifugation, followed by two washes with ice-cold 1x PBS; pellets were re-suspended in chilled 70% ethanol and kept at 4°C. For cell cycle analysis, cells were pelleted by centrifugation, resuspended in 400μl PI (propidium iodide)/RNAse Staining Solution (Cell Signaling, MA; 4087S) with 0.1% Tween and incubated for one hour, protected from light. After the incubation, cells were spun down and washed twice with 1x PBS and resuspended in 500μl of 1x PBS for analysis by flow-cytometry. Flow cytometry was performed using the CytoFlex (Beckman Coulter; Brea, CA, USA) and results were provided by the flow cytometry core facility at Stony Brook University and visualized via Modfit.

## RESULTS

### Expression of the *SGMS1* TSS-7 transcript is regulated by two upstream regions in its promoter

The first step in understanding regulation of *SGMS1* gene expression was to identify regions in the TSS-7 promoter that are critical for its activity in the K562 CML cell line. Progressive 5’ deletion constructs of the full-length TSS-7 promoter (sequence in **Supplementary Table 1)** were generated, and promoter activity was measured using the luciferase reporter assay **(Figure 1A)**. Deletion of 5’ bases up to -160 bp did not significantly affect TSS-7 promoter activity as compared to the full-length sequence while deletion of bases between -160 bp to -127 bp resulted in a 50% reduction in promoter activity. Further deletions of the promoter did not impact activity until deletion of bases between -118 bp to -115 bp when promoter activity was almost completely lost (93%). Additional downstream deletions had no further effect. Thus, region I (-160 to -127) and region II (-118 to -115) **(Figure 1B)** contribute significantly to the promoter activity upstream of TSS-7.

**Figure 1.**
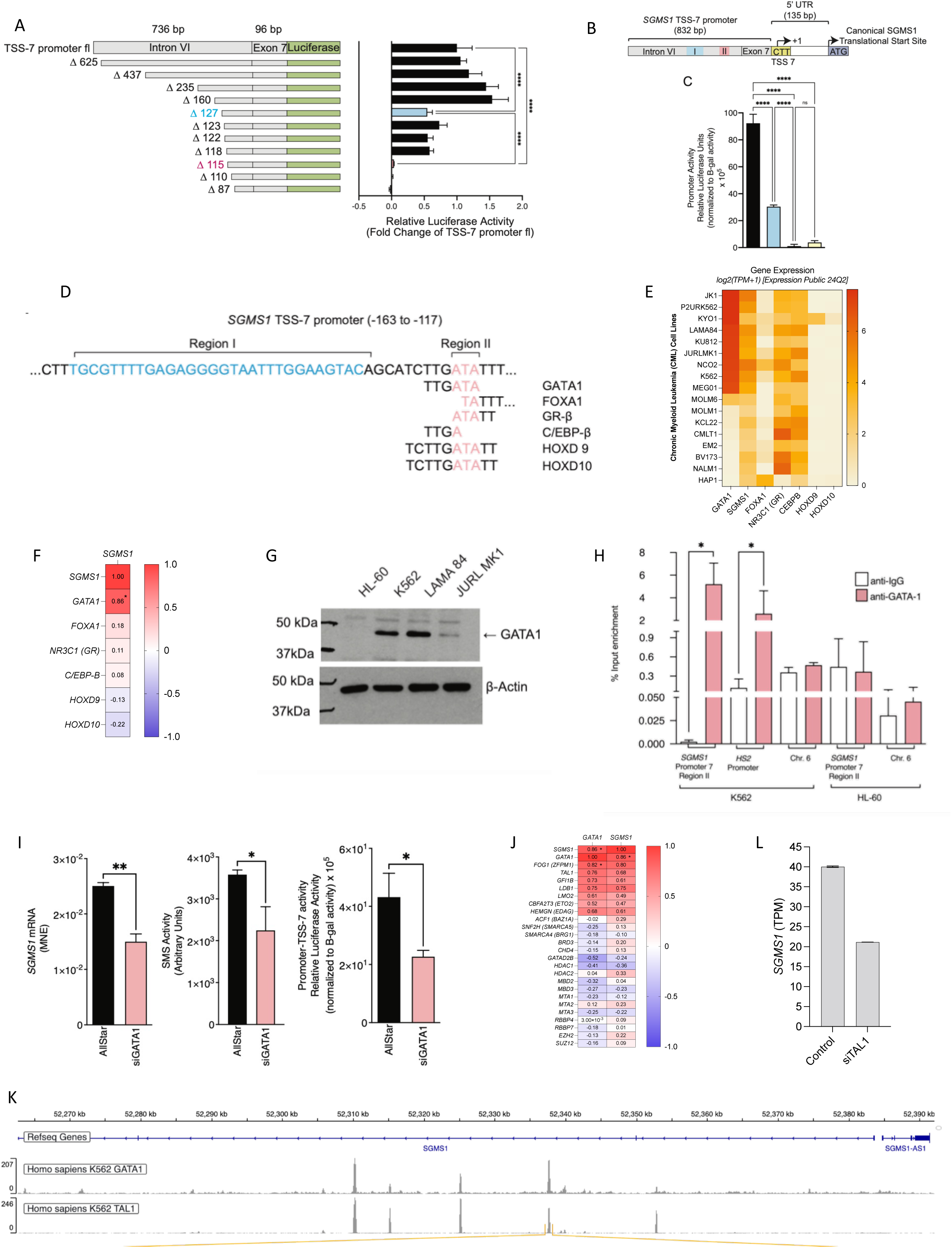
*SGMS1* is under the direct control of the GATA1-TAL1 transcriptional complex. **A: (Left panel)** Schematic representation of the 5’ deletion constructs of *SGMS1* TSS-7 promoter. Grey boxes indicate the TSS-7 promoter region labeled with canonical exon/intron annotations. The green box indicates the luciferase gene under the regulation of these promoter constructs. The construct names indicate the remaining number of bases of the TSS-7 full-length promoter (TSS-7 promoter fl). **(Right panel)** Promoter activity of the different *SGMS1* TSS-7 promoter 5’ deletion constructs in luciferase activity assay. Luciferase activity was measured as previously described (34). For each replicate, luciferase activity was normalized to β-galactosidase (β-gal) activity (relative luciferase units - RLU), subtracted from the normalized luciferase activity (RLU) of the empty vector, and expressed as fold change compared to RLU of the full-length TSS-7 promoter. The results represent 3 independent experiments. Significance calculated with One-way ANOVA with post-hoc Tukey’s multiple comparison test; **** p < 0.0001. **B:** Schematic representation of the two *cis*-regulatory regions identified by promoter deletion constructs analysis. Grey regions depict the TSS-7 promoter and its span within the *SGMS1* locus. Blue box represents Region I, pink/red box indicates Region II. White box depicts the transcribed region downstream of the transcriptional start site TSS-7(yellow box); translational start site is depicted by the purple box. **C:** Relative luciferase activity was calculated on the deletion of individual regions and compared to that of the full-length TSS-7 promoter. Results represent 3 independent experiments. Significance calculated with One-way ANOVA with post-hoc Tukey’s multiple comparison test; **** p < 0.0001. **D:** Figure shows the base-sequence map of Region I (blue) and Region II (red) on the *SGMS1* TSS-7 promoter (and some flanking sequences). Below the base-sequence map of Region I and II are the predicted consensus binding motifs for transcription factors identified by PROMO analysis with sites spanning Region II. Highlighted in red/pink are bases with sequence similarity to Region II for each transcription factor consensus sequence. **E:** Heatmap depicts the mRNA expression of the six putative transcription factors mapping to Region II of the *SGMS1* TSS-7 promoter. mRNA expression for the putative transcription factors was downloaded from DepMap (Expression Public 24Q2) and plotted into a heatmap on Graph Pad Prism. **F:** Pearson’s correlation of mRNA expression of the six transcription factors with *SGMS1* using DepMap data base. **G:** Representative western blot for GATA1 and β-actin expression in control cell line HL-60 and CML cell lines K562, LAMA-84 and JURL-MK-1. Lysates were probed for protein expression of GATA1 and β-actin (as loading control). Full gel image is presented in **Supplementary Figure S1B** for reference. **H:** Chromatin-immunoprecipitation (ChIP) was performed in K562 and HL-60 cells using IgG (white) and GATA1 (red) antibodies. Quantification of GATA1-occupied DNA was performed by qRT-PCR; primers were designed to target the genomic Region II of *SGMS1*-TSS 7, GATA1 binding site in the promoter of *HS2* (positive control) and chromosome 6 (negative control). Y-axis measures the % input enrichment. Results represent 3 independent experiments (except for Chromosome 6 which is representative of 2 independent experiments). Asterisks indicate significance; Unpaired t-test *p<0.05. **I:** Endogenous *SGMS1* mRNA expression was quantified by qRT-PCR in K562 cells after *GATA1* levels were reduced with siRNA for 24h. The first panel shows mean of normalized expression (MNE) of *SGMS1* calculated after normalization to β-actin. Results are from 3 independent experiments. **p < 0.005. The second panel shows endogenous SMS activity which was measured in K562 cells in which *GATA1* had been downregulated by siRNA. The figure shows the endogenous total normalized SMS activity (expressed as arbitrary units). Results are from 3 independent experiments. *p< 0.05. The third panel shows *GATA1* downregulation on TSS-7 promoter activity in K562 cells. Relative luciferase activity of TSS-7 (full-length) promoter is shown in cells transfected with Control (AllStar siRNA) and siGATA1 (siRNA for *GATA1*). Results are from 3 independent experiments. *p < 0.05. **J:** Pearson’s correlation matrix of mRNA expression of the GATA1 partners with *GATA1* and *SGMS1* in CML cell lines. mRNA expression was derived from the CCLE RNAseq data and correlation was calculated in DepMap Data Explorer 2.0 using the Pearson correlation coefficient. **K:** GATA1 and TAL1 ChIP-seq profile within the *SGMS1* locus in K562 cells show co-localization. **L**: *TAL1* downregulation results in reduced *SGMS1* Transcripts Per Kilobase Million (TPM) based on Encode datasets ENCSR641CMW (control); ENCFF714HMY (control); ENCFF035JUJ (siTAL1); and ENCSR336ZWX (siTAL1).

The individual contributions of these two regions on promoter activity were assessed using deletion constructs of either region I, or region II, or both region I and II. Promoter activity for the three constructs was quantified **(Figure 1C)**. Deletion of region I alone resulted in a 50% loss in promoter activity, while loss of region II alone or region I and region II resulted in almost complete loss of promoter activity, mirroring the loss in activity seen with the 5’ deletion construct activity analysis. These results suggest that the TSS-7 promoter has two *cis*-regulatory regions important for regulating transcription from TSS-7, region I which potentially cooperates with region II, the core promoter.

### GATA1 binds to the core promoter region of *SGMS1* TSS-7 promoter

Sequence analysis of region II, the core promoter of TSS-7, was conducted with PROMO (40, 41) (http://alggen.lsi.upc.es/cgi-bin/promo_v3/promo/promoinit.cgi?dirDB=TF_8.3) to identify potential transcription factor binding sites (TFBSs). Analysis revealed six putative DNA binding motifs for transcription factors (TFs) GATA1, FOXA1, GR-β (NR3C1), C/EBP-β, HOXD9 and HOXD10 **(Figure 1D)**.

We assessed expression of the cognate TFs in K562 cells and additional CML cell lines known to have high *SGMS1* expression (34) by evaluating mRNA expression data from the Cancer Cell Line Encyclopedia (CCLE) (42–44) (**Figure 1E).** *GATA1* had the highest expression in the CML cells followed by *GR-β* (NR3C1), *C/EBP-β* and *FOXA1,* while *HOXD9* and *HOXD10* had negligible expression. Next, we determined the correlation between expression of these TFs relative to *SGMS1* across these cell lines **(Figure 1F).** By far, the strongest positive significant correlation was between *GATA1* and *SGMS1*. High GATA1 protein levels were confirmed by western-blotting in CML cell lines known to have high *SGMS1* expression (34) compared to HL-60 cells (as a negative control) **(Figure 1G, Supplementary Figure 1).** These results indicate that GATA1 and SGMS1 are highly expressed in several CML cell lines and their expression is highly correlated.

To determine if GATA1 binds to region II, Chromatin Immunoprecipitation (ChIP) was performed with GATA1 antibody or non-targeting control IgG antibody in K562 and GATA1-negative HL-60 cells. qRT-PCR was used to quantify occupancy at three genomic regions: the *SGMS1* region II sequence encompassing the putative GATA1 DNA binding motif, the HS2 region of the β-globin locus control region known to be GATA1-occupied (45–47) as positive control, and a third region within chromosome 6 [primer sequence from (48)] which lacks known GATA1 motifs within the amplified region +/-200 bps. The ChIP analysis **(Figure 1H)** revealed GATA1 occupancy at *SGMS1* region II and β-globin HS2 in K562 cells. As expected, the region in chromosome 6 (negative control) was not recovered. Furthermore, no enrichments were detected in HL-60 cells (which lack GATA1), supporting specific binding of GATA1 to the *SGMS1* region II locus in K562 cells.

### GATA1 and TAL1 regulate *SGMS1* expression

To further analyze the importance of GATA1 in regulating *SGMS1* transcription, we downregulated *GATA1* by siRNA in K562 cells and quantified *SGMS1* mRNA expression by qRT-PCR **(Figure 1I, Supplementary Figure 2A-B)**. *GATA1* downregulation significantly decreased *SGMS1* mRNA compared to cells transfected with control siRNA (**Figure 1I, first panel**). The extent of the changes was comparable to the effect on *ALAS2* (**Supplementary Figure 2C**), an established GATA1 target (46, 49, 50). As K562 cells express both *GATA1* and *GATA2* (51), we tested whether downregulating *GATA2* influenced *SGMS1* expression, but it did not (**Supplementary Figure 3**). Furthermore, we asked whether reduced *SGMS1* expression upon *GATA1* downregulation impacted SMS enzymatic activity and found that total *in vitro* SMS activity (SMS1+SMS2) was significantly reduced (**Figure 1I, second panel).**

Next, we probed whether GATA1 regulates *SGMS1* TSS-7 promoter activity. Following *GATA1* downregulation, cells were co-transfected with the *SGMS1* TSS-7 promoter-luciferase plasmid or empty vector and the beta-galactosidase expression construct. Luciferase activity was quantified after 16 hours **(Figure 1I, third panel)**. Downregulation of *GATA1* significantly inhibited activity of the full-length TSS-7 promoter construct, suggesting that GATA1 regulates *SGMS1* promoter activity. Together, these results indicate that GATA1 regulates TSS-7 promoter activity of *SGMS1* and increases *SGMS1* expression.

GATA1 functions as both activator and repressor of gene transcription based on the factors it complexes with (52). Well known GATA1 partners include TAL1, LDB1, ACF1, SNF2h, Gfi-1b, FOG1, and MeCP1(52–54). To identify which binding partner might work in complex with GATA1 to regulate *SGMS1* transcription, we took a bioinformatics approach and identified the degree of correlation for each partner with *SGMS1* in CML cell lines using the CCLE expression data **(Figure 1J**). TAL1 and LDB1 (two binding partners of GATA1 which work in concert to promote erythroid differentiation)(55, 56) had strong positive correlations, as did Gfi-1b, which works with GATA1 to repress proliferation in hematopoietic stem cells, while its overexpression promotes cell proliferation in M6 and M7 AMLs (57–59). To investigate regulation of *SGMS1* by *TAL1,* GATA1 and TAL1 ChIP-seq profile at the *SGMS1* locus was determined in K562 cells using publicly available datasets and found that several peaks for GATA1 and TAL1 co-localized in Intron II (**Figure 1K**). Both canonical and non-canonical DNA binding consensus sequences for GATA1 (^A^/_T_ GATA ^G^/_A_; NGATA^G^/_C_; NGATGG)(60) and TAL1 (CAGNTG; ATGGT)(61, 62) were identified in correspondence of the peaks, and are characterized in **Supplementary Table 6**. ChIP-seq data relative to other GATA1 binding partners were not available. Analysis of RNA-seq data in K562 cells from the Encode Project found that downregulation of *TAL1* reduces *SGMS1* Transcripts Per Kilobase Million (TPM) by half (**Figure 1L**). Altogether, these results support the direct positive regulation of *SGMS1* by GATA1 in complex with TAL1.

### *SGMS1* is regulated by GATA1 in erythroleukemic (M6) and megakaryocytic (M7) Acute Myeloid Leukemia (AML) cell lines

As GATA1 expression has been reported in other leukemic cell types (63), we extended the expression analysis to incorporate all leukemia cell lines, as well as *TAL1*, and explore whether GATA1 targeting of *SGMS1* might have broader relevance. The majority of CML cell lines (labeled in pink in Figure 2A) demonstrated a high expression of *GATA1* and, in agreement with results in Figure 1, there was a positive correlation with *SGMS1*. Remarkably, a strong positive *GATA1*/*TAL1*/*SGMS1* correlation was also observed in a defined subset of Acute Myeloid Leukemia (AML) cell lines (TF-1, MO7E, UT7, MOLM-16, Hel, HEL92.1.7, CMK, CMK115, CMK 86, F36P and OCIM1); these cells belong exclusively to the M6 or M7 AML subtypes (**Figure 2A and Supplementary Figure 4A**). High GATA1 expression was confirmed in MOLM-16 and Hel cells by western blotting, and importantly, this was tracked by high expression (**Figure 2B**) and enzymatic activity (**Figure 2C**) of SGMS1. Knockdown of *GATA1* by siRNA in Hel cells resulted in 50% down-regulation of *SGMS1* quantified by qRT-PCR (**Figure 2D**). Importantly, in *GATA1* positive AML patient samples, *TAL1* showed a high correlation coefficient (0.67) and was the best positively correlated GATA1 partner with *SGMS1* (**Supplementary Figure 4B)**.

**Figure 2.**
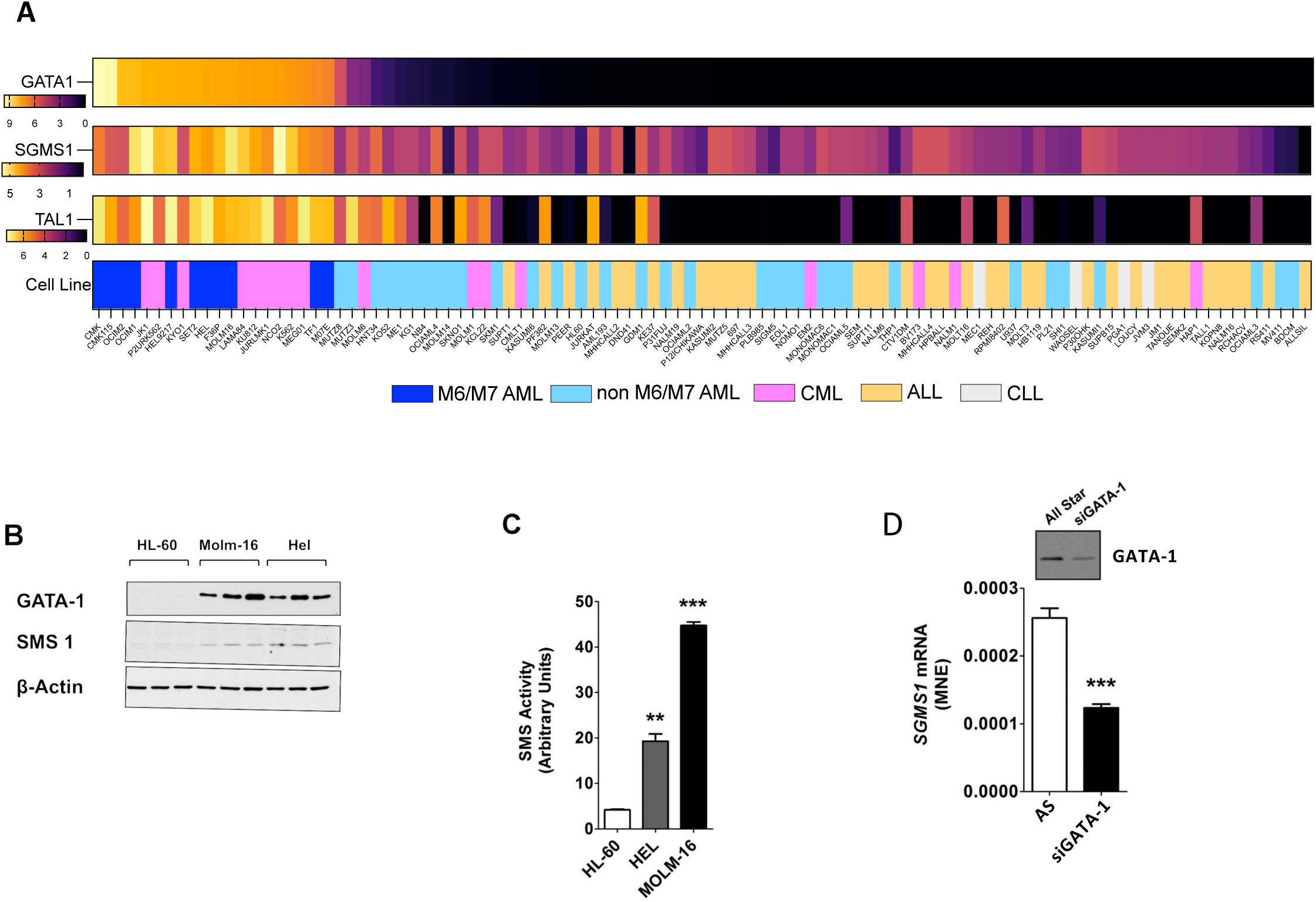
*SGMS1* is regulated by GATA1 in Acute Myeloid Leukemia. **A**: Heat-map showing the relative mRNA expression of *GATA1*, *SGMS1,* and *TAL1* as derived from CCLE. The expression is shown across an array of leukemic cell lines indicated below the heat-map and color coded based on their leukemia type. **B**: Protein expression of GATA1 and SMS1 protein was evaluated in AML cell lines HEL, MOLM-16 and HL-60. Cell lysates were extracted from these cell lines and blotted using gene specific antibodies. β-Actin protein blotted as loading control. The blot shows samples from three independent experiments. **C**: Total SMS activity from cell lysates was measured in HL-60, MOLM-16 and HEL cells. The SMS activity shown in figure is derived from band-quantification of fluorescent lipid product from a TLC plate and normalized to lysate protein. Values are the mean and standard deviation from three independent experiments. **D** siRNA down regulation of *GATA-1* in HEL was assessed by western blotting 48h post-siRNA transfection of AllStar Control (AS) or siRNA for *GATA-1* (siGATA-1). This figure is representative of two independent experiments. (**Right panel**) Endogenous *SGMS1* expression was measured by qRT-PCR in HEL cells in which GATA-1 expression had been down-regulated by siRNA. The figure shows mean of normalized expression (MNE) of *SGMS1* calculated after normalization to expression of β-Actin. Values are the mean and standard deviation from three independent experiments. (Asterisks indicate significance; **** p<0.00005; *** p<0.0005; ** p<0.005).

Altogether, these results support a role for GATA1/TAL1 in regulation of *SGMS1* in M6 and M7 AMLs.

### *SGMS1* regulates cell-cycle progression at the G2/M checkpoint

Considering the highly aggressive nature of M6/M7 AML and the urgent need for effective treatments, we set to study the functional implications of the newly discovered elevated SGMS1 activity using Hel cells as a model for M6 AML. *SGMS1* expression was down-regulated in Hel cells for 6 days with transfections occurring every other day to ensure maximal reduction. After the third and final transfection, cells were collected for gene expression analysis by microarray. The efficiency of *SGMS1* down-regulation was quantified by RT-PCR (**Figure 3A)** and showed over 50% *SGMS1* knock-down; this was confirmed by using the Transcriptome Analysis Console to evaluate the microarray results. Gene-Set Enrichment Analysis (GSEA) revealed that *SGMS1* downregulation induced the most significant changes in gene sets associated with G2/M checkpoint and mitotic spindle (**Figure 3B-D**); gene expression associated with protein secretion and E2F targets was also significantly affected. These results suggested that SGMS1 function may intersect with processes involved with cell proliferation, especially around mitosis.

**Figure 3:**
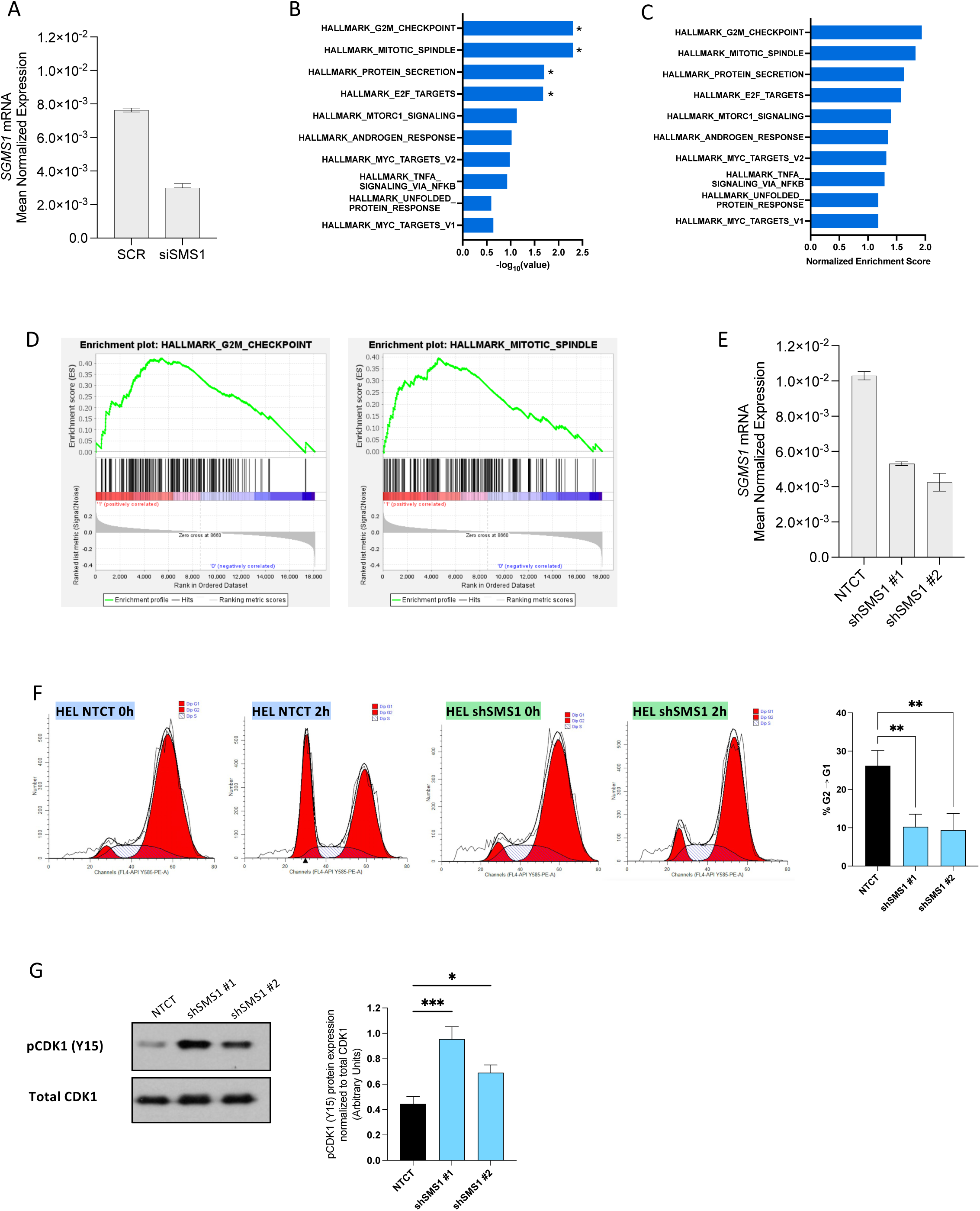
*SGMS1* regulates cell-cycle progression at the G2/M checkpoint. **A-D**: Transfection of Hel cells with *SGMS1* or SCR (control) siRNA (total of 3 transfections, one transfection every other day for six days). Results from 4 independent experiments. **A:** *SGMS1* expression by qRT-PCR; beta actin used as normalization control to calculate the Mean of Normalized Expression. **B-D:** Analysis of gene expression changes as measured by microarray after *SGMS1* or SCR (control) siRNA in Hel cells. **B:** The false discovery rate q-values; * indicates an adjusted p value <0.05. **C:** Normalized enrichment scores for gene sets enriched in cells with *SGMS1* downregulation. **D:** Enrichment plots of HALLMARK_G2M_CHECKPOINT and HALLMARK_MITOTIC_SPINDLE gene signatures found in siSGMS1 versus SCR control. **E:** *SGMS1* mRNA measured by qRT-PCR in cells with short hairpin(sh)-RNA mediated-stable downregulation of *SGMS1* (Stable shRNA clones - NTCT: Non targeting control; shSGMS1: *SGMS1* stably downregulated). **F:** Representative cell cycle plots of NTCT and shSGMS1#1 Hel cells at zero hours and two hours after release from nocodazole-mediated G2/M arrest. Bar graph representation of the percentage of cells that transitioned from G2 to G1-phase two hours post release from G2/M arrest ** p<0.005. **G: Left panel**, western blot showing protein levels of phosphorylated CDK1 at tyrosine 15 (pCDK1 Y15), and total CDK1 in control sh-RNA cells (NTCT) and in cells with stable *SGMS1* downregulation (shSMS1, two clones). **Right panel**, quantification of pCDK1 protein levels normalized to total CDK1 protein.

To assess the biological relevance of the most significantly enriched gene-sets - G2/M checkpoint and mitotic spindle- a stable knock-down of *SGMS1* in Hel cells with shRNA was first established, single clones isolated, and the two clones with the strongest *SGMS1* knockdown were selected (**Figure 3E)**. We then set out to assess whether *SGMS1* knockdown functionally impacted the G2/M checkpoint by specifically evaluating its effect on re-entry into G1 after arrest in G2/M with nocodazole (64). After the G2/M arrest, cells were washed and collected immediately (0h) and at two hours after wash (**Figure 3F**). At two hours post-release, cells with *SGMS1* knockdown were significantly slower to resume normal cycling, with a 60% reduction in cells going from G2 to G1 compared to the non-targeting control (**Figure 3F)**. The delayed exit from the G2/M arrest correlated with increased levels of phosphorylated CDK1 protein (**Figure 3G**); CDK1 is a catalytic subunit of the M phase-promoting factor (MPF) complex, and its phosphorylation renders it inactive, hindering mitotic progression (65–67). Together these results support a role for SGMS1 in regulating exit from G2/M in M6 AML cells.

### High *SGMS1* expression in AML patients is associated with worse probability of survival and SGMS1 inhibition sensitizes M6 AML Hel cells to anti-mitotic treatment

In line with the observations in M6 and M7 AML cell lines, AML patients of the erythroid and megakaryocytic subtypes (M6 and M7 AML, respectively) have higher *SGMS1* expression compared to other subtypes and healthy subjects as observed in clinical samples (**Figure 4A**). High *SGMS1* expression is also associated with significantly decreased survival probability of AML (**Figure 4B**). Therefore, we tested whether inhibition of SGMS1 would be detrimental to M6 AML Hel cells, particularly in the presence of anti-mitotic drugs given SGMS1’s function in promoting G2/M transition. Notably, downregulation of *SGMS1* sensitized Hel cells to the microtubule stabilizer and mitotic inhibitor Taxol (68–70), as measured by percent of trypan blue positive cells (**Figure 4C**). To confirm involvement of the SM synthetic node of sphingolipid metabolism in this phenotype, we tested the effect of HPA-12, an inhibitor of the ceramide transfer protein CERT1, which provides ceramide to SGMS1 for SM synthesis. Like knock down of *SGMS1*, HPA-12 effectively sensitized Hel cells to Taxol (**Figure 4D**). As inhibition of CERT1 has been linked to sensitization of chemotherapy *via* induction of ER stress specifically by activation of the PERK-mediated arm (71), we determined changes in phospho-PERK upon inhibition of SGMS1 and in the presence of Taxol (**Figure 4E**). As shown in the blot, down-regulation of *SGMS1* caused a significant increase in phosphorylation of PERK in the presence of Taxol, indicative of activation of PERK and enhanced ER stress.

**Figure 4:**
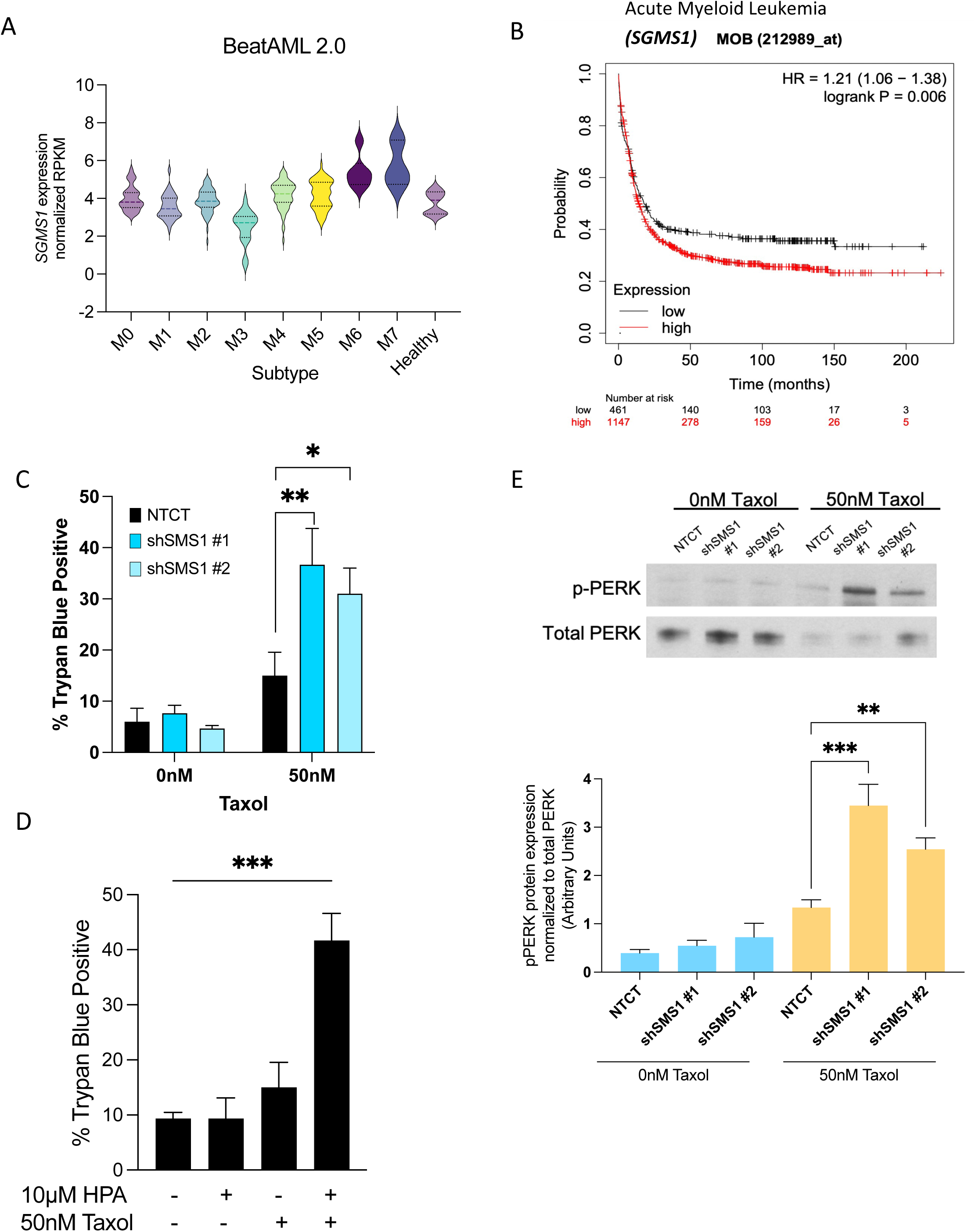
High *SGMS1* expression is associated with poorer survival probability in AML and counteracts cell cycle targeting therapy. **A:** *SGMS1* gene expression level in AML patients (BeatAML2.0) as reported in Vizome stratified by AML subtype and including healthy clinical samples. **B:** Kaplan-Meier survival curve for high and low *SGMS1* expression in AML patients visualized via KM plotter (93). **C:** Percentage of dead cells (trypan blue positive) in cells with stable downregulation of *SGMS1* (two clones: shSMS1 #1 and shSMS1 #2) compared with non-targeting control (NTCT) with or without 50nM Taxol. **D:** Cytotoxicity in Hel cells after 48 hours treatment with Vehicle, 10µM HPA, 50nM Taxol, or a combination. Cell death is measured as percentage of trypan blue positive cells. **E:** (**Top panel**) Western blot showing protein levels of phosphorylated PERK (p-PERK) and total PERK with and without Taxol treatment (0nM or 50nM) in control cells (NTCT) and in cells with stable *SGMS1* downregulation (shSMS1, two clones). (**Bottom panel**) Quantification of p-PERK levels normalized to total PERK protein. * p<0.05, ** p<0.01, ***p<0.005

Altogether these results support a role for SGMS1 in ensuring optimal progression through G2/M and they point to inhibition of SGMS1/SM synthesis as a potential novel strategy for sensitization of M6 AML cells against anti-mitotic agents like Taxol.

## DISCUSSION

We have previously demonstrated that *SGMS1* is transcriptionally upregulated in K562 cells via utilization of its alternative transcription start site TSS-7 (34). In this work, we characterized the novel TSS-7 promoter and identified GATA1 as a transcriptional regulator of *SGMS1* in a complex with TAL1. The link among GATA1/TAL1/high SGMS1 exists in all GATA1 positive leukemic cells, including M6/M7 AML. Functionally, high *SGMS1* regulated exit of M6 AML cells from G2/M and its down-regulation contributed to sensitization towards the anti-mitotic agent, Taxol. This role for SGMS1 may provide a novel strategy to sensitize M6 (and possibly M7) AML cells to chemotherapy with anti-mitotic agents.

First, the *SGMS1* TSS-7 promoter mediates transcription of *SGMS1* mRNA encoding the full protein with an enhanced efficiency of translation (34, 36). Notably, this promoter is unique, marked by the absence of conventional (but not exclusive) features seen in alternative promoters such as TATA-box, BRE elements and/or Inr sequences (72, 73). While the TSS-7 promoter does not contain these elements, it included two important regulatory regions, with region I contributing to promoter activity and region II functioning as the core promoter in a transfection assay. Deletion of region II abrogated TSS-7 promoter activity and, while encompassing a non-canonical GATA1 binding DNA motif, region II was GATA1-occupied. In K562 cells, GATA1, but not GATA2, elevated *SGMS1* expression; and ChIP analysis confirmed GATA1 occupancy at region II of the *SGMS1* TSS-7 promoter. GATA1 expression correlated with *SGMS1* across CML and M6 and M7 AML cells, suggesting that GATA1-mediated transcriptional regulation of *SGMS1,* validated in loss-of-function assays, is broadly relevant. While we demonstrated that GATA1 occupies TSS-7 promoter in K562 cells, ChIP-seq analysis revealed GATA1 occupancy within intron II, which contains canonical and non-canonical GATA1 motifs. The molecular bases of this discrepancy are being investigated.

Additionally, our analysis supports a role for TAL1 together with GATA1 in regulating *SGMS1* expression. GATA1 typically works in complex with different partners that can either repress or activate expression of target genes. One such binding partner is TAL1, which plays roles in maintaining multipotency of hematopoietic stem cells, and in leukemogenesis(74). Interestingly, high expression of TAL1 can also lead to malignant transformation of hematopoietic cells(75). In our analysis of publicly available ChiPSeq data, we find that *TAL1* colocalized with GATA1 in intron II of *SGMS1*, that *GATA1*/*TAL1*/*SGMS1* expression strongly correlated in GATA1 positive leukemias (cell lines and patient samples) and that TAL1 knock-down in K562 cells significantly reduced *SGMS1* expression. Using the correlation between *GATA1/TAL1* and *SGMS1* uncovered in CML cells, we were able to define another subset of leukemic cells that displays this same pattern - M6 and M7 AML cells. The M6/M7 AML model is interesting because in addition to being leukemic cells with high GATA1/high SGMS1 and therefore suitable for the study of SGMS1 functions, M6/M7 AML cells are very aggressive and do not respond to standard of care and patients with M6/M7 subtypes have the worst prognosis across all AMLs (76, 77). Therefore, new findings in these cells might also provide the basis for new therapeutic concepts to be tested.

The induction of GATA1 in normal hematopoietic progenitors is essential for erythropoiesis (50, 52, 78, 79). On the other hand, high GATA1 level in most CML cell lines and all M6/M7 AML cells is not sufficient to induce erythroid differentiation and it might even sustain proliferation and promote chemotherapeutic resistance as it was shown in blasts from patients with acute megakaryocytic leukemia treated with cytarabine and daunorubicin (80). Interestingly, when we compared GATA1 gene targets in fetal peripheral blood-derived erythroblasts (PBDEs) to those in erythroleukemic K562 cells, notable differences were found (*manuscript submitted for publication*). While in fetal PBDE the top two most significant pathways targeted by GATA1 involved heme biosynthesis and elimination of organelles (processes that are crucial for terminal erythroid differentiation) (81, 82), in K562 erythroleukemic cells, the top hits were pathways of sphingolipid signaling and sphingolipid metabolism (including the *SGMS1* gene). This suggests that the gene expression response to GATA1 in erythroleukemia seems to be different from normal erythroblasts and sphingolipid genes are a prime GATA1 target in erythroleukemia. As resistance to chemotherapies remains a major challenge for M6 and M7 AML patients(83), targeting sphingolipid enzymes (SGMS1 and/or others) in conjunction with anti-mitotic drugs may represent a novel strategy to improve the chemotherapeutic response of these aggressive and difficult to treat AMLs.

In GATA1^+^ M6 AML Hel cells, we uncovered a role for *SGMS1* in regulating exit from G2/M, as confirmed by GSEA analysis of microarray data and functional assays (cell cycle analysis, changes in check point marker proteins and sensitivity to the anti-mitotic drug, Taxol). Interestingly, CERT1 has also been implicated in chemotherapy resistance (including Taxol) (71, 84). SGMS1 and CERT1 functions are linked, in that CERT1 is responsible for transporting ceramide to the trans-Golgi (85) where SGMS1 converts it into SM. This points to the possibility that synthesis of SM (*via* CERT1 and SGMS1) may ultimately contribute to microtubule integrity and G2/M transition, conceivably implicating the lipids regulated by these proteins, ceramide, SM and/or DAG. Also, both downregulation of *SGMS1* and inhibition of CERT1 were found to increase activation of PERK (**Figure 4** and (71, 84)); as ceramide accumulation has been placed both upstream and downstream of phospho-PERK (86–90) and very limited information links SM or DAG to phospho-PERK, the mechanistic link between inhibition of SM synthesis and activation of PERK is not clear at this time. The fact that blockade of CERT1 sensitizes cells to Taxol similar to knockdown of *SGMS1* is also relevant from a pharmacological/therapeutic point of view. While specific SGMS1 inhibitors validated in cell studies have not been reported, inhibitors of CERT1 are available, albeit at micromolar active concentrations. However, given the recent link established between hyperactivation of CERT1 and neurological abnormalities (91, 92), it is predicted that additional efforts towards the identification of specific and potent CERT1 inhibitors will be forthcoming. This would provide more effective tools to target SM synthesis which could then be also tested for sensitization of M6/M7 AML cells.

## Data Sharing Statement

Requests for further information on reagents, methodologies and data should be directed and will be met by.

## Acknowledgements

The authors wish to acknowledge the expert support of the Genomics Core Facility at Stony Brook University for microarray analysis and the Flow Cytometry Core Facility at Stony Brook University for assistance with flow cytometry. Additionally, the authors wish to recognize the critical assessment of the manuscript by Dr. Yusuf A. Hannun and expert input by Dr. Emery Bresnick. This work was supported by U.S. National Institutes of Health, National Cancer Institute Grant P01 CA097132 (to CL for Project #4), and pilot funding from Stony Brook Cancer Center/The John C. Dunphy Private Foundation to CL. Y.R. is a 2023-2024 American Fellow of the American Association for University Women and a recipient of a Turner Dissertation Fellowship by the Stony Brook University’s Center for Inclusive Education.

## Authorship Contributions

Y.R. proposed the hypotheses, designed and performed the experiments, analyzed the data, prepared figures and wrote the manuscript; S.M. proposed the hypotheses, designed and performed the experiments, analyzed the data, prepared figures and edited the manuscript; G-Q.Y. performed experiments; S.B.C. analyzed the data, prepared figures and edited the manuscript; C.V.S. analyzed the data; C.L. proposed the hypothesis, designed the study, analyzed the data, prepared figures and wrote the manuscript.

