## Supplemental Figure legends, Tables and Figures for "Regulation of cell cycle by the novel GATA1/TAL1/Sphingomyelin Synthase 1 (*SGMS1*) transcriptional axis. Implications for anti-leukemic strategies"

**SUPPLEMENTARY INFORMATION**

**Supplementary Figure Legends**

**Supplementary Figure 1.**

**A.** Quantification of GATA1 protein level normalized to β-actin in the control cell line HL-60 and CML cell lines K562, LAMA-84 and JURL-MK-1. Results represent 3 independent experiments. **B.** Representative full western blot gel image. Top panel shows GATA1 expression and bottom panel shows corresponding β-actin expression.

**Supplementary Figure 2.**

**A.** siRNA-mediated downregulation of GATA1 in K562 cells was assessed by western blotting. GATA1 antibodies were used to measure abundance of GATA1 protein expression at 24h post-siRNA transfection of Control (AllStar negative control from Qiagen) or siGATA1 (siRNA for *GATA1*). β-actin protein blotted as loading control. This figure is representative of three independent experiments. **B.** Quantification of GATA1 protein levels normalized to β-actin in K562 cells after down-regulation of GATA1 by siRNA at 24 hours. Results are from 3 independent experiments. **$p < 0.002$. **C.** Endogenous *ALAS2* mRNA expression was measured by qRT-PCR in K562 cells in which *GATA1* expression was reduced by siRNA for 24h. The figure shows Mean of Normalized Expression (MNE) of *ALAS2* calculated after normalization to β-actin. Results are from 3 independent experiments. ****$p < 0.0001$.

**Supplementary Figure 3.**

**A.** siRNA-mediated down regulation of GATA-2 in K562 cells was assessed by western blotting. GATA-2 antibodies were used to measure the extent of down-regulation in GATA-2 expression 48h post-siRNA transfection of Control (AllStar) or siGATA-2 (siRNA for GATA-2). B-Actin protein blotted as loading control. Results are from 2 independent experiments. **B.** Endogenous *SGMS1* mRNA expression was quantified by qRT-PCR in K562 cells after *GATA2* levels were reduced with siRNA for 48h. The first panel shows mean of normalized expression (MNE) of *SGMS1* calculated after normalization to β-actin.

**Supplementary Figure 4.** Correlation coefficients for GATA1 binding partners in AML cell lines and GATA1 positive AML patient samples.

**A:** Pearson's correlation matrix of mRNA expression of the GATA1 partners with *GATA1* and *SGMS1* in AML cell lines. mRNA expression was derived from the CCLE RNAseq data and correlation was calculated in DepMap Data Explorer 2.0 using the Pearson correlation coefficient. **B:** TAL1 shows the highest positive correlation coefficient among all GATA1 partners with *SGMS1* in GATA1 positive AML patient samples. Data are derived from the LAML dataset from The Cancer Genome Atlas (TCGA).*denotes components of the MeCP1 transcriptional complex.

#### Supplementary Tables:

##### Supplementary Table 1: TSS-7 promoter sequence.

Canonical *SGMS1* Intron VI
Canonical *SGMS1* Exon 7

```
TTCTGTGAGTTGCTGTATGTTATGTACATTTTTAAACATTCTTCACGAGGTAGAGTAAGCAAATATATTTTATTCTTTTAAATTATCTCAGGAAGCATGTAAGGACAAAGAG
AATACCTGGTCTCTTGACCAGGTTGAAAAGTGTCCAAAGCACTGTCAAGTTGCTTCTTAGAAAATAGACCATTGGAAATTATTATTATTTTTTTTTTTTTGAGAAGGAATCTC
GCTCTGTCGCCCAGGCTGGAGTACAGTGGCTGATCTCGGCTCACTGCAAGCTCCGCCTCCCGGGTTCACACCATTCTCCTGCCTCAGCCTGGGCAGGAGTAGCTGGGACTAC
AGGCGCCCGCCACCACACCCGGCTAATTTTTTGTATTTTTAGTAGAGACAGGGTTTCACCGTGTTAGCCAGGATGCTCTCTATCTCCTGACCTCGTGATCCGCCCGCCTCGT
TCTCCCAAAGCGCTGGGATTACAGGCGTGAGCCACCACGCCCGGCTGACCGTTGGAAATTCTATAAAATATGTCTCCTTTGACAACATCCATTTTCAGATCCAATTCAATGT
GTGTCGCAAAAGACTAATAACCAGGAAATCAGTAGTCCCTGAAACGAATACATGTTTGAAAACTATTAAGTTTACCAATAACTGGTTTTCTACTTTCCTTATGTATATAAGG
CTTTGCGTTTTGAGAGGGGTAATTTGGAAGTACAGCATCTTGATATTTTCTCTTGCTTTCACAGAAGGAAGAATCCTGCTCAAAAATGAGGTGAATTAATACTTGGGCGCTC
AGGAACCCTGGACAGCTACATGAGGTGTTTAAAAACTGCCCTGACCAT
```

##### Supplementary Table 2: Primer sequences for TSS-7 5' progressive promoter deletion constructs.

Each forward primer name indicates the final truncated length of the TSS-7 promoter generated.

| Primer | Sequence (5' to 3') |
|---|---|
| Δ 625 | TTGAGAAGGAATCTCGCTCTGTCG |
| Δ 437 | CGTGTTAGCCAGGATGCTCT |
| Δ 235 | CCTGAAACGAATACATGTTTGAA |
| Δ 160 | TGCGTTTTGAGAGGGGTAAT |
| Δ 127 | AGCATCTTGATATTTTCTCTTGCTTTC |
| Δ 123 | TCTTGATATTTTCTCTTGCTTTCA |
| Δ 122 | CTTGATATTTTCTCTTGCTTTCACAGA |
| Δ 118 | ATATTTTCTCTTGCTTTCACAG |
| Δ 115 | TTTTCTCTTGCTTTCACAGAAGG |
| Δ 110 | TCTTGCTTTCACAGAAGGAAGAATC |
| Δ 87 | TCCTGCTCAAAAATGAGGTG |
| Promoter 7-reverse | ATGGTCAGGGCAGTTTTTAAA |

##### Supplementary Table 3: TSS-7 cis-regulatory region deletion construct sequences.

Δ Region I:

```
TAAGCAGGTACCTCTGTGAGTTGCTGTATGTTATGTACATTTTTAAACATTCTTCACGAGGTAGAGTAAGCAAATATATTTTATTCTTTTAAATTATCTCAGGAAGCATGTA
AGGACAAAGAGAATACCTGGTCTCTTGACCAGGTTGAAAAGTGTCCAAAGCACTGTCAAGTTGCTTCTTAGAAAATAGACCATTGGAAATTATTATTATTTTTTTTTTTTTGA
GAAGGAATCTCGCTCTGTCGCCCAGGCTGGAGTACAGTGGCTGATCTCGGCTCACTGCAAGCTCCGCCTCCCGGGTTCACACCATTCTCCTGCCTCAGCCTGGGCAGGAGTA
GCTGGGACTACAGGCGCCCGCCACCACACCCGGCTAATTTTTTGTATTTTTAGTAGAGACAGGGTTTCACCGTGTTAGCCAGGATGCTCTCTATCTCCTGACCTCGTGATCC
GCCCGCCTCGTTCTCCCAAAGCGCTGGGATTACAGGCGTGAGCCACCACGCCCGGCTGACCGTTGGAAATTCTATAAAATATGTCTCCTTTGACAACATCCATTTTCAGATC
CAATTCAATGTGTGTCGCAAAAGACTAATAACCAGGAAATCAGTAGTCCCTGAAACGAATACATGTTTGAAAACTATTAAGTTTACCAATAACTGGTTTTCTACTTTCCTTA
TGTATATAAGGCTTAGCATCTTGATATTTTCTCTTGCTTTCACAGAAGGAAGAATCCTGCTCAAAAATGAGGTGAATTAATACTTGGGCGCTCAGGAACCCTGGACAGCTAC
ATGAGGTGTTTAAAAACTGCCCTGACCATCTCGAGTAAGCA
```

Δ Region II:

```
TAAGCAGGTACCTCTGTGAGTTGCTGTATGTTATGTACATTTTTAAACATTCTTCACGAGGTAGAGTAAGCAAATATATTTTATTCTTTTAAATTATCTCAGGAAGCATGTA
AGGACAAAGAGAATACCTGGTCTCTTGACCAGGTTGAAAAGTGTCCAAAGCACTGTCAAGTTGCTTCTTAGAAAATAGACCATTGGAAATTATTATTATTTTTTTTTTTTTGA
GAAGGAATCTCGCTCTGTCGCCCAGGCTGGAGTACAGTGGCTGATCTCGGCTCACTGCAAGCTCCGCCTCCCGGGTTCACACCATTCTCCTGCCTCAGCCTGGGCAGGAGTA
GCTGGGACTACAGGCGCCCGCCACCACACCCGGCTAATTTTTTGTATTTTTAGTAGAGACAGGGTTTCACCGTGTTAGCCAGGATGCTCTCTATCTCCTGACCTCGTGATCC
GCCCGCCTCGTTCTCCCAAAGCGCTGGGATTACAGGCGTGAGCCACCACGCCCGGCTGACCGTTGGAAATTCTATAAAATATGTCTCCTTTGACAACATCCATTTTCAGATC
CAATTCAATGTGTGTCGCAAAAGACTAATAACCAGGAAATCAGTAGTCCCTGAAACGAATACATGTTTGAAAACTATTAAGTTTACCAATAACTGGTTTTCTACTTTCCTTA
TGTATATAAGGCTTTGCGTTTTGAGAGGGGTAATTTGGAAGTACAGCATCTTGATTTTCTCTTGCTTTCACAGAAGGAAGAATCCTGCTCAAAAATGAGGTGAATTAATACT
TGGGCGCTCAGGAACCCTGGACAGCTACATGAGGTGTTTAAAAACTGCCCTGACCATCTCGAGTAAGCA
```

Δ Region I and II:

```
TAAGCAGGTACCTCTGTGAGTTGCTGTATGTTATGTACATTTTTAAACATTCTTCACGAGGTAGAGTAAGCAAATATATTTTATTCTTTTAAATTATCTCAGGAAGCATGTA
AGGACAAAGAGAATACCTGGTCTCTTGACCAGGTTGAAAAGTGTCCAAAGCACTGTCAAGTTGCTTCTTAGAAAATAGACCATTGGAAATTATTATTATTTTTTTTTTTTGA
GAAGGAATCTCGCTCTGTCGCCCAGGCTGGAGTACAGTGGCTGATCTCGGCTCACTGCAAGCTCCGCCTCCCGGGTTCACACCATTCTCCTGCCTCAGCCTGGGCAGGAGTA
GCTGGGACTACAGGCGCCCGCCACCACACCCGGCTAATTTTTTGTATTTTTAGTAGAGACAGGGTTTCACCGTGTTAGCCAGGATGCTCTCTATCTCCTGACCTCGTGATCC
GCCCGCCTCGTTCTCCCAAAGCGCTGGGATTACAGGCGTGAGCCACCACGCCCGGCTGACCGTTGGAAATTCTATAAAATATGTCTCCTTTGACAACATCCATTTTCAGATC
CAATTCAATGTGTGTCGCAAAAGACTAATAACCAGGAAATCAGTAGTCCCTGAAACGAATACATGTTTGAAAACTATTAAGTTTACCAATAACTGGTTTTCTACTTTCCTTA
TGTATATAAGGCTTAGCATCTTGATTTTCTCTTGCTTTCACAGAAGGAAGAATCCTGCTCAAAAATGAGGTGAATTAATACTTGGGCGCTCAGGAACCCTGGACAGCTACAT
GAGGTGTTTAAAAACTGCCCTGACCATCTCGAGTAAGCA
```

#### Supplementary Table 4: Primary and secondary antibodies for Western Blotting.

| **Primary antibodies** | | | |
|---|---|---|---|
| **Protein** | **Description, catalogue number** | **Dilution** | **Company** |
| GATA1 | N1 rat monoclonal, sc-266 | 1:5000 | Santa Cruz Biotechnology, Dallas, TX |
| GATA2 | H-116 rabbit polyclonal, sc-9008 | 1:5000 | Santa Cruz Biotechnology, Dallas, TX |
| Actin | I-19 goat polyclonal, sc-1616 | 1:2500 | Santa Cruz Biotechnology, Dallas, TX |
| SMS1 | X1702P rabbit polyclonal against full length protein | 1:1000 | ExAlpha, Shirley, MA (33, 34, 36) |
| **Secondary antibodies** | | | |
| goat anti-rat | Cat # 112-035-003 | 1:5000 | Jackson ImmunoResearch Laboratories, West Grove, PA |
| goat anti-rabbit | Cat # 111-035-003 | 1:5000 | Jackson ImmunoResearch Laboratories, West Grove, PA |
| rabbit anti-goat | Cat # 305-035-003 | 1:5000 | Jackson ImmunoResearch Laboratories, West Grove, PA |

#### Supplementary Table 5: qRT-PCR primers for measuring *SGMS1* (34) and *ALAS2* (94) mRNA.

| **Gene** | **Sequence (5' to 3')** | |
|---|---|---|
| Exon 7 *SGMS1* | Forward | GCCAGGACTTGATCAACCTAACC |
| | Reverse | CCATTGGCATGGCCGTTCTTG |
| *ALAS2* | Forward | CAGTTCCTGTTTGGTATTGGACG |
| | Reverse | TGCCTTCTGCACAATCTTGCT |
| *ACTB* | Forward | ATTGGCAATGAGCGGTTCC |
| | Reverse | GGTAGTTTCGTGGATGCCACA |
| *18S RNA* | Forward | CTCAACACGGGAAACCTCAC |
| | Reverse | CGCTCCACCAACTAAGAACG |

**Supplementary Table 6: GATA1 (bolded) and TAL1 (italicized) canonical and noncanonical binding sequences found within peaks on intron II of *SGMS1***

| location | sequence |
| --- | --- |
| 52,310,100 bp – 52,310,450 bp | CAAAGGAGGTTGACGAGGAGGGGTCAGAGAAG*CAGGTG*TATTAGTACAATCTCAGGGCATGCCAGGGTAGCAGATTGGTGAGTGTGCCTTTTTAAGTCAAGGTTCCG**GGATAGATAG**TATAGGAGTCTGGCTGCTGGAGAGGGCTGGGAGGAAGGGCAGTAGA**AGATAA**GGCTGGAGCCAGAGGTGTTGGCCTTGAAAACCAGTCCAAGGCATGTTAACTCTCTTTGGCAGGAAGGGTGTATTGTAGGGC**TGATAA**AAGCTGTGCTTCA**GGATAG**CTAGTCCGCAGCTTCATGCAAATT**AGATAG**AGGGATTAGAACTTG*CAGTTG*CTTCATTGGGAAATTTGGAAATGGGTGCTTGGTGAGGTAATGAAAAGAGGCAGGAAGCAGTGGCCTATTATTTGC |
| 52,315,100 bp – 52,315,350 bp | ATTAACATTTTAAATAGGAAGTTTACAGTTCTTCCCTCTTTTAATAAATGTTTCCTTGTATAAGAATGATCTTTGAAGCAGAGGCCTCAACCGCATTCCTTATTACACTGCTGCAGCTCTGGATGTACCCATGAAGCCCTGTAATTGGAATCTTCTGGCTGGCTC**AGATAA**GCAGCTTATTTGTGAGCCAGGTCCTGCCCTGTTTCTTTAAGGCCTCAAAGCAGGAAG*ATGGT*GAACTTTATGCAATTCTGAGTTGACTCCAATC*CAGATG*TATTTTGGAACTAGGTAGAATATGACTGG |
| 52,325,050 bp – 52,325,350 bp | TAGAGTTGAGATTTTAACTCATTGTGCTTGACTTCCAAACCCATGCCCTTAACCACTTCTTTCCAGACGTATCTTTTGTGTAACATAA*ATGGT*GGCATTGACAAATGCTTTCCCAATCTCCACCCCAAAGGGTGCTGCTTACTTTCTGTTCCAAGC**AGATAC**AGTAGAAGTCAGTACTGCCCTGATGTAGATTACATTGTCATCTTTTTTACAGCAGCAGGAAGTTCTTCAGTGGCAGTATAGATGACACATGTGATTTTGTATAAAAATATATATTTTT |
| 52,337,500 bp – 52,337,850 | GGTCTCTGCAAAAATTAAGCAGTTTTAGAAGAGCAGAGTGGTTCTTTTACTATAGGCTAACTGACCTTGGATTATTTCATGATCTGCTCCTTGAGG**TGATAA**GCT*CAGTTG*CTTCTTGTCTTTTCTGAACGTTACATGTATCCTCTGTCAAATACCAGTCCTTCCTTCTAGCTTACTGGAACACTTAGTCATTGTGTAGTAACCAAC*ATGGT*CATTTCTGGCCCTGGCGCTGATT**AGATAA**CAGAAGTCCCCTTCACAAGCTTCGCAGCTCTGTGCAGTAGCAGCCCTTTAATAGCTGTAACTTCACCACTGGGAAAGGAAAGGAAAGGAAAGCCTCTCTGCTTTCATGCTGATTGCTATGGAAGCTTGCCCA**AGATAC**TTCCCCTTGCATGTGCAGGGCTGTGTGC |
| 52,352,650 bp – 52,352,970 bp | GATCTGGTTCCA**AGATGG**CTCACTCACATAGCTGTTGCCAGGAGGCCTCAGTTCCTTGCTGGTTCTTGGCAAGAGGCCTCAGTTCTTTCCCACGTGTATCTCTGCAGAGCTTCTTGAGCGTGTCCTCATATCAGAACAGCCAGCTTCCCTCCGAAAACAG**AGATGG**AAGGTGGAGTCTTTGGACTTTTGCAGTCTCACGCCAGCATTTCAGCAATATTGTAGTGGTTCCCACAT |

### Supplementary Figure S1

A B

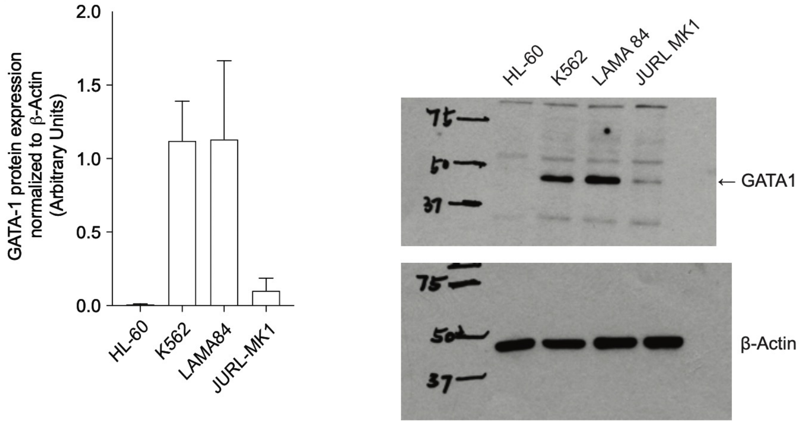

### Supplementary Figure S2

A B C

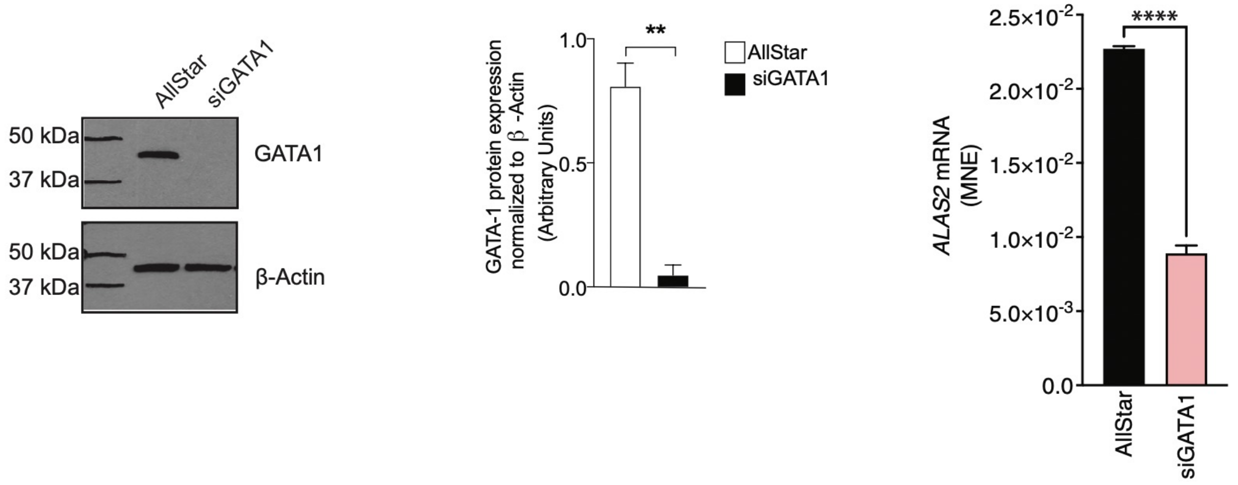

### Supplementary Figure S3

**A** **B**

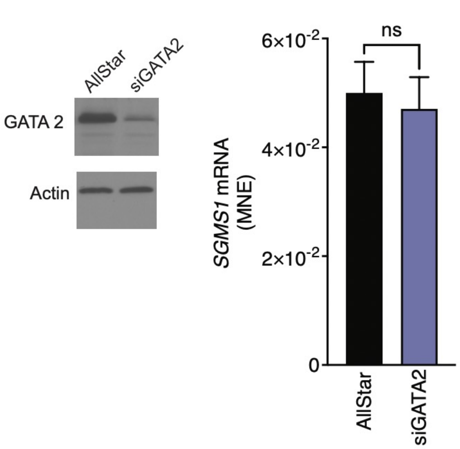

### Supplementary Figure S4

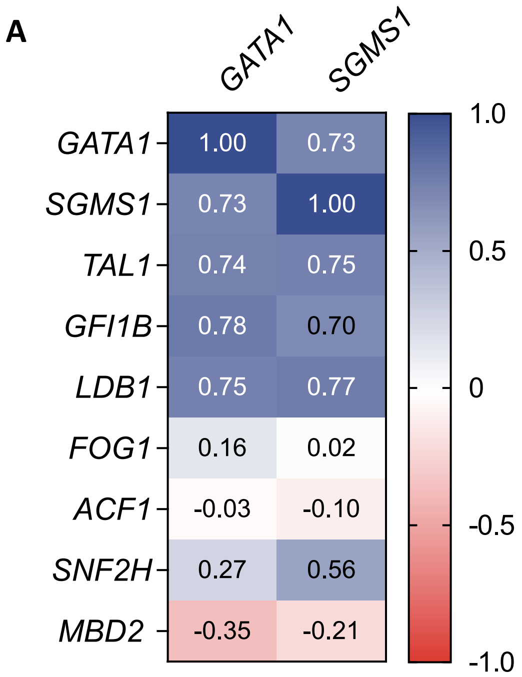

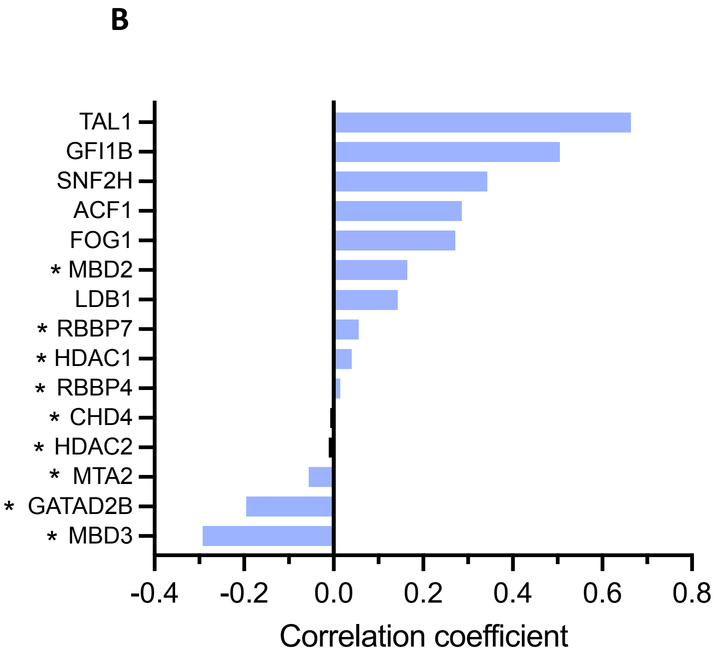
